# Multiscale assessment of oviposition habitat associations and implications for management in the spotted lanternfly (*Lycorma delicatula*), an emerging invasive pest

**DOI:** 10.1101/2022.09.08.507131

**Authors:** Victoria A. Ramirez, Sebastiano De Bona, Matthew R. Helmus, Jocelyn E. Behm

**Affiliations:** Integrative Ecology Lab, Temple University, 1925 N. 12^th^ Street, Philadelphia, PA USA

**Keywords:** *Acer*, *Ailanthus altissima*, Control strategies, Habitat selection, Habitat use, Hierarchical spatial scale, Multiscale management

## Abstract

1. Control of incipient invaders—established invasive species in the early stages of spreading— can be inhibited by incomplete knowledge of the species’ habitat use. By identifying consistent habitat associations for incipient invaders early, control efforts can be more effective. Yet, because habitat associations are the result of multiscale processes, approaches are needed for integrating data collected across scales to identify them.
2. We employed a hierarchical, multiscale approach to identify oviposition habitat associations in the spotted lanternfly (*Lycorma delicatula*) an incipient invasive species of high concern in the United States. We targeted the oviposition habitat spatial scales most likely to be used by lanternflies and the spatial scales of explanatory environmental variables most easily used by managers to locate egg masses to control. We explored whether habitat associations exist for oviposition habitat use and how well habitat use is explained by the duration sites have been invaded. Finally, because habitat associations are likely driven by fitness, we determined if the use of any habitat types were associated with higher fitness as measured by the number of eggs per egg mass (i.e., fecundity).
3. Spotted lanternflies exhibited oviposition habitat associations at the landscape, site, and tree scales. Overall, lanternflies oviposited more frequently at sites and on trees with low canopy cover in the surrounding landscape, on trees from the *Acer* genus, and in the crowns of larger trees beyond the reach of managers without special equipment. The duration a site had been invaded had opposing effects at the site and tree scales which indicates the need for scale-dependent management approaches.
4. Despite the consistent habitat associations identified, no environmental variables explained variation in lanternfly fecundity, indicating more work is needed to identify environmental drivers of spotted lanternfly fitness.
5. *Synthesis and applications*. Our results indicate a multiscale approach is needed for spotted lanternfly control with unique strategies for locating egg masses at sites and on trees that vary in invasion duration. Additionally, our results suggest that multiscale investigations of habitat associations would likely inform the control of other invasive species as well.

## Introduction

Because invasive species are being introduced at accelerating rates (Seebens et al., 2017) and are a significant cause of negative ecological and economic impacts (Pyšek et al., 2020) devising strategies for their control is imperative. Incipient invaders – established species in the early stages of spreading – are thought to be the most cost-effective invasion stage to control (Homans & Horie, 2011), yet their control still provides unique challenges. Since incipient invaders are often at low densities and managers have incomplete knowledge of the species’ habitat use, searches for them can be long, ineffective, and costly (Mehta et al., 2007). However, if incipient invaders are consistently associated with particular habitat types, the early identification of their habitat associations can inform management by drastically reducing search efforts and improving success.

In general, *habitat use* refers to the aspects of the habitat species exploit for their needs (Krausman, 1999) and *habitat associations* arise when species consistently use the same type of habitat, often because that habitat provides higher fitness (McLoughlin et al., 2006). For incipient invaders to exhibit habitat associations driven by fitness, they must reliably identify cues that indicate habitat quality in a novel environment. Successful invasive species likely interpret habitat quality cues correctly (e.g., Rondoni et al., 2017), but the extent to which habitat associations are driven by fitness for incipient invaders has not been well-explored. When incipient invaders are generalist species and/or experience enemy release, many habitat types could yield high fitness. Instead, fitness may be limited by their own population density; when sites are initially colonized and populations are at low density, fitness may differ significantly from later stages when densities are higher (Greene & Stamps, 2001). In this case, although species’ distributions are driven by fitness, consistent habitat associations may be absent, and species’ distributions may be better explained by the duration sites have been invaded than by habitat variables. Finally, habitat associations may arise when incipient invaders are more abundant in habitats associated with dispersal. For example, cavity-nesting mosquitoes are often found nesting in tires in human settlements in their invaded range because they are directly transported in tires to these locations by humans (Lounibos, 2002). Consequently, habitat associations due to dispersal may not be directly related to fitness. Exploring the relationship between habitat associations and fitness may provide useful insight into how incipient invaders are using and benefitting from their newly invaded range.

Regardless of the mechanisms generating them, the processes that underlie habitat associations operate at multiple spatial scales (Mayor et al., 2009). Investigations that employ a hierarchical design where habitat use at multiple, nested spatial scales is explained by habitat variables, provide a more complete understanding of habitat associations than investigating single or non-nested scales (Johnson, 1980). In this hierarchical context, habitat associations may arise between habitat use and environmental variables acting at the same scale and/or broader scales. From a management perspective, hierarchical designs provide useful guidance for enacting management actions for invasive species (Martins et al., 2016). Broad-scale habitat associations indicate the general area where a species may be, while fine-scale associations provide key information for exactly where to find the species to control. Therefore, it is important to conduct multiscale assessments at both the scales at which species use habitat and the scales at which management decisions are made (Brown & Barney, 2021). Despite their utility, multiscale assessments of habitat associations for non-native species are scarce (Biggs & Olden, 2011; Froehly et al., 2020), partly because they may require different skillsets to collect finer scale data through field studies and broader scale data aggregated through data science methods. Studies that can integrate datasets across scales are poised to provide comprehensive information to direct management efforts.

Here, we conduct a multiscale assessment of habitat associations in the spotted lanternfly (*Lycorma delicatula*), an incipient invasive species of high concern in the US (Dara et al., 2015). Native to China, the spotted lanternfly planthopper was first documented in the US in Pennsylvania in 2014 and as of 2021 has expanded its range to nine additional states in the mid-Atlantic region (NYSIPM, 2021). Its reproduction is univoltine; females lay wax-covered egg masses on trees in the autumn, which hatch into nymphs the following spring and then transition to adults in late summer (Liu, 2019). Control efforts currently target adult, nymph, and egg stages, yet because egg masses can be difficult to detect, identifying oviposition habitat associations would help to inform management and control efforts substantially.

Evidence suggests that spotted lanternflies may have oviposition habitat associations driven by fitness and/or dispersal, or alternatively, oviposition habitat use may be better explained the duration a site is invaded. Although spotted lanternfly has been documented using over 150 plant taxa (Barringer & Ciafré, 2020), their fitness varies significantly according to host plant species (Murman et al., 2020; Uyi et al., 2020, 2021). Most of the host plant species that provide high fitness for spotted lanternflies, like *Ailanthus altissima*, are weedy or ornamental species that are common in human-dominated habitats. Furthermore, experiments indicate that spotted lanternflies respond to host plant volatile cues (Derstine et al., 2020) indicating that spotted lanternflies may be able to select habitats that provide high fitness. Additionally, spotted lanternfly disperse with human assistance within their invaded range (Urban et al., 2021) which may further bias habitat associations toward human-dominated habitats. Finally, spotted lanternfly population dynamics likely follow a boom-and-bust pattern where they colonize a site at low density, densities increase over time and then drop again. This indicates the duration a site has been colonized may explain habitat use in addition to or better than habitat variables.

We use a hierarchical design to identify environmental variables influencing multiscale oviposition habitat use from the landscape to egg mass scale (Fig. 1). In addition, we use fecundity as a proxy for fitness to determine if fitness is associated with any habitat types. We integrate data science methods with traditional field methods to generate explanatory environmental variables at two spatial scales likely to influence spotted lanternfly habitat associations in addition to being easily identifiable and usable by managers. Finally, we discuss how the multiscale nature of our results has clear implications for spotted lanternfly control and management.

**Figure 1.**
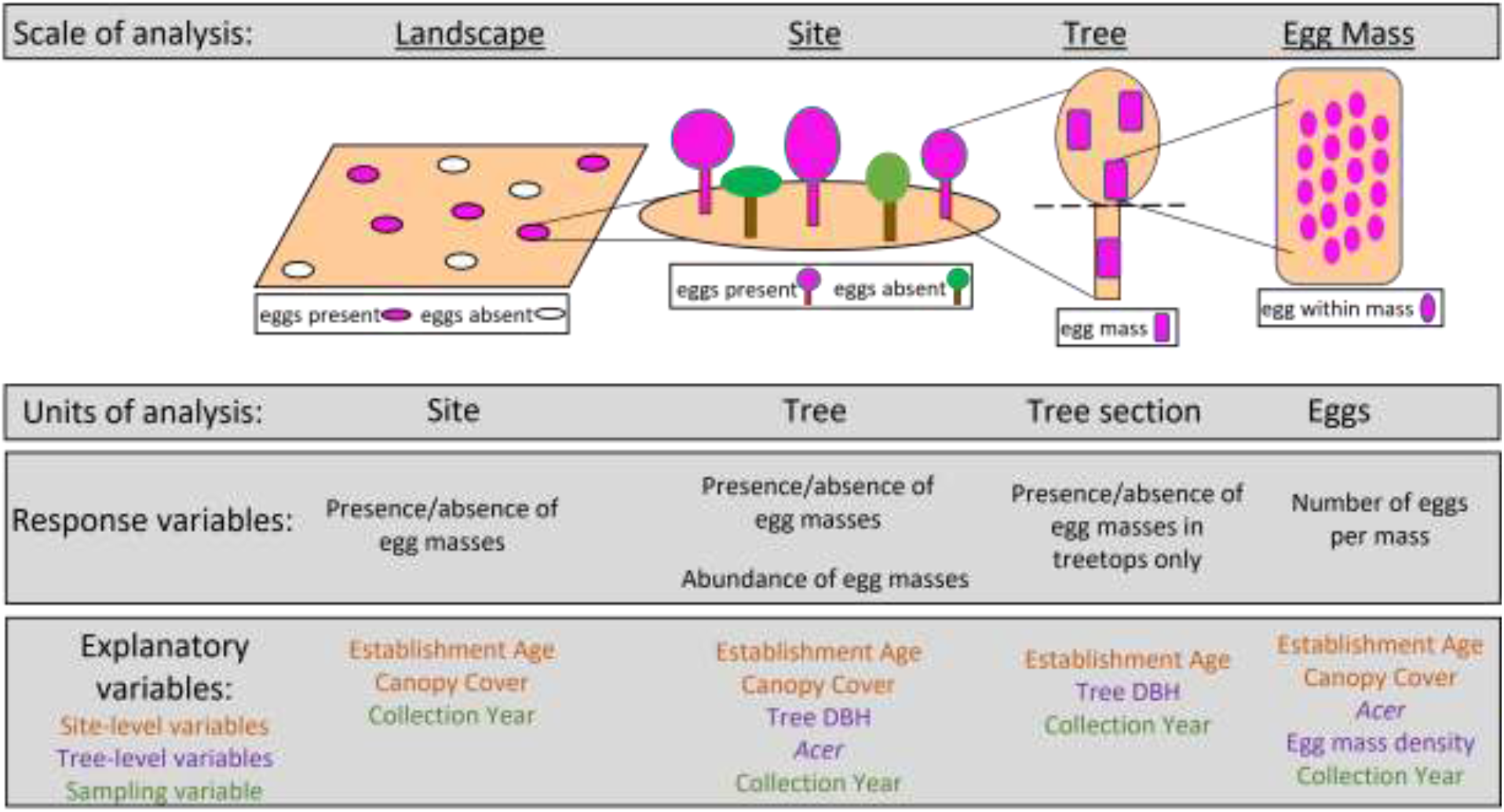
Overview of hierarchical study design showing the four scales of oviposition habitat use response variables, the unit of analysis, and the site- and tree-level explanatory variables used in our analyses.

## Methods

We analyzed spotted lanternfly oviposition habitat use and fecundity across four hierarchical scales: oviposition at sites across Pennsylvania (landscape-scale), oviposition on trees within a site (site-scale), oviposition within a tree (tree-scale), and eggs within an egg mass (i.e., fecundity; egg mass scale) (Fig. 1). We use fecundity both as a scale of habitat use and a proxy for fitness to determine what habitats provide higher fitness. For clarity, we use the term ‘scale’ to refer to the different scales of these response variables of spotted lanternfly habitat use, and the term ‘level’ to refer to the scales of explanatory variables (McGarigal et al., 2016).

Our oviposition and fecundity response variables for the four scales came from two surveys. The first was a large, multiagency, statewide survey of egg masses at 141,984 sites across Pennsylvania (hereafter ‘statewide survey’; Fig. 2A) and we used these data for the landscape-scale analysis. For the finer three scales, we used data from an in-depth survey of 66 sites sampled from the “core” (longest invaded sites) to the “edge” (newer invaded sites) of the invaded range in southeastern Pennsylvania (hereafter ‘core-to-edge survey’; Fig. 2B).

**Figure 2.**
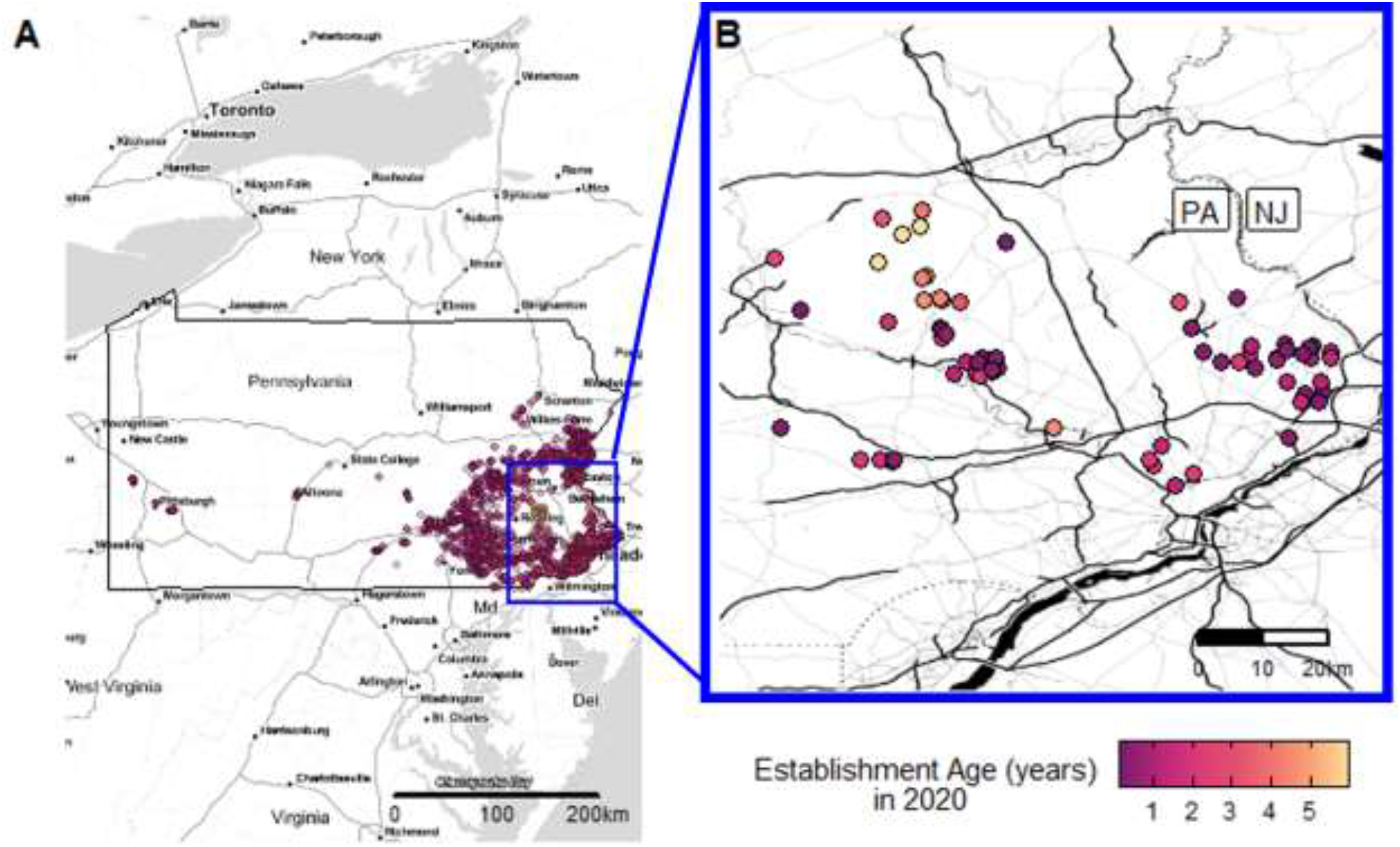
Multiscale oviposition habitat selection by the invasive spotted lanternfly was assessed using data from two surveys: (A) A statewide survey recorded the presence/absence of eggs at 7,363 sites across Pennsylvania. (B) A core-to-edge survey recorded egg mass abundance and eggs per mass on multiple trees within sites. Sites sampled in both surveys varied in the number of years spotted lanternfly was established at sites (establishment age – site shading).

Below we describe our data collection and multiscale analyses on these two datasets. Explanatory variables were scaled to facilitate comparison of their estimated effect sizes among models at each scale. Analyses were conducted using R programming for statistics version 4.1.1 (R Core Team, 2021). Model diagnostics were run for each model using the “simulateResiduals” function from the DHARMa package (Hartig, 2020) and collinearity among variables (variance-inflation) was checked using the “vif” function from the car package (Fox & Weisberg, 2019).

### Statewide survey

To assess oviposition habitat use at the landscape scale, we used egg mass presence/absence at sites within the invaded range across Pennsylvania from surveys by the Pennsylvania Department of Agriculture (PDA) and the United States Department of Agriculture (USDA). At each site, these agencies recorded the presence/absence of spotted lanternflies and egg masses during the 2018-19 and 2019-20 seasons (hereafter ‘collection year 2018’ and ‘collection year 2019’, respectively) across Pennsylvania. Although 141,984 sites were surveyed, many sites had no spotted lanternfly present because the agencies surveyed beyond the edge of the invasion range. To restrict the dataset to only sites within the invaded range, we defined the invaded range by creating 0.1 km grids across the state and analyzed only those sites contained in grid cells where at least one site had an established spotted lanternfly population. This grid size corresponds to the known distance that spotted lanternfly adults disperse during the mating and egg laying season (Wolfin et al., 2019). This filtering resulted in 7,363 sites within 4,016 grid cells that contained at least one site with an established lanternfly population.

### Core-to-edge survey

We sampled egg masses at 66 sites spanning the invasion core to edge (Fig. 2B) in southeastern Pennsylvania for two reproductive seasons: November 20, 2018 to May 17, 2019 (hereafter ‘collection year 2018’) and November 14, 2019 to March 9, 2020 (hereafter ‘collection year 2019’). Across the two collection years, 24 sites were sampled during both years, 28 sampled in 2018, and 14 sampled in 2019. At each site, we recorded oviposition and fecundity at three scales: site, tree, and egg mass (Fig. 1). At the site scale, we counted the number of egg masses per tree, aiming for a minimum of five trees per site. At the tree scale, we recorded if egg masses on a tree were only in the treetops or distributed across the trunk and treetops. At the egg mass scale, we collected at least five egg masses per tree and counted the number of eggs per egg mass with a dissecting microscope. The number of eggs per egg mass was our metric of fecundity and the response variable in the egg mass-scale analysis. All trees in the survey were identified to species and DBH (cm) measured.

### Derived site-level explanatory variables

To test if spotted lanternfly oviposition is associated with human-dominated habitats, we calculated canopy cover in 500m buffers surrounding each site from both surveys. Lower canopy cover habitat may convey higher fitness due to the presence of the spotted lanternfly’s preferred host plant species, and/or dispersal of lanternfly to new sites. Canopy cover was the average pixel value within the buffer of the Global Forest Cover Change (GFCC) Tree Cover dataset computed using Google Earth Engine at 30m resolution (Sexton et al., 2013).

To test if spotted lanternfly oviposition and fecundity are influenced by establishment age—the duration a site has had an established population of lanternfly—we estimated the establishment age of each site. To do this, we spatially interpolated presence/absence of spotted lanternfly at any life stage from data collected by PDA and USDA across PA for each year between 2015 and 2019 (Supporting Information 1, section 1). From our interpolation method, the minimum value for any site was zero and the maximum value was 5 years, indicating certainty of spotted lanternfly being established throughout the 5 years considered, between 2015-2019 (Fig. 2). Using this method, we calculated establishment ages for sites from both the statewide and core-to-edge surveys.

### Landscape scale analysis

For our landscape scale analysis, we explored habitat associations across sites in the statewide survey. We tested the effect of surrounding canopy cover, establishment age, and collection year on egg mass presence at sites with a logistic regression (“glm” in the stats package). We included collection year to account for unexplained year-to-year variation that may influence oviposition and accounted for spatial autocorrelation among sites using the “autocov_dist” function from the spdep package (Bivand, 2022; Dormann et al., 2007).

### Site scale analysis

For our site-scale analysis, we explored which variables affected oviposition across trees within the sites sampled in our core-to-edge survey. We included the site-level establishment age and canopy cover variables due to their possible association with oviposition on trees. Since tree size and tree taxonomy may influence oviposition habitat use (Liu, 2019; Liu & Hartlieb, 2020), we included tree DBH and tree species as tree-level variables. However, our dataset encompassed 32 tree species from 23 genera, which was more categories than we had statistical power to test. Therefore, we grouped trees into two categories for which spotted lanternfly likely did or did not exhibit an oviposition preference at the genus level. Specifically, we calculated spotted lanternfly oviposition preference as the selection of a genus for oviposition (proportion of egg masses per genus at a site) relative to the availability of oviposition substrate provided by that genus at a site (proportion of total DBH per genus at a site) (Beyer et al., 2010). We found spotted lanternfly oviposited on the genus *Acer* at a higher frequency than its availability at sites (Supporting Information 1, section 2). Accordingly, we grouped all genera into *Acer/*not *Acer* categories and use this as our binary tree taxonomy variable in subsequent analyses to test the strength of the preference for tree taxonomy relative to other environmental variables influencing habitat use.

Our models for the site-scale analysis to explain variation in oviposition on trees within sites included the explanatory variables: establishment age, canopy cover, collection year, tree size (DBH) and tree taxonomy (*Acer*/ not *Acer*). Given the high number of trees we sampled with zero egg masses, we conducted our site-scale analysis in two parts. First, we tested the effect of the explanatory variables on the presence/absence of egg masses on trees using a binomial generalized linear mixed model (GLMM) with site as a random effect (“glmer” in lme4 [Bates et al., 2015]). Only data from year 2019 were used because trees without eggs were not recorded in collection year 2018. Then, using only the subset of the trees in our dataset that exhibited oviposition activity (i.e. excluding trees with zero egg masses), we fit a second model testing the effect of the explanatory variables on the number of egg masses per tree using a GLMM with a zero-truncated negative binomial error distribution with a *log-link* function (`glmmTMB` [Brooks et al., 2017]) using data from both collection years.

### Tree scale analysis

For our tree-scale analysis, we explored the effect of the explanatory variables on the distribution of egg masses within a tree because lanternflies may prefer to oviposit in treetops over trunks (Liu & Hartlieb, 2020). Our binary response variable was the presence of egg masses only in the treetops (1) versus presence of egg masses in the treetops and along the trunk (0). We never observed instances of egg masses only on the trunk and not in the treetops when multiple masses were present. This analysis only included trees for which egg masses were present. To explore these patterns, we fit a GLMM with a binomial error distribution and a *logit-link* function, with establishment age, collection year, and tree DBH as the fixed effects and site as a random effect (“glmer” in `lme4`).

### Egg mass scale analysis

For our egg mass scale analysis, we focused on explanatory variables that could influence fecundity: establishment age, canopy cover, collection year, tree taxonomy (*Acer*/not *Acer*), and egg mass density. Egg mas density per tree was included in the model because planthopper density may be negatively correlated with fecundity (Heong 1988; Denno and Roderick 1992). Egg mass density was calculated as the number of egg masses per tree scaled by tree DBH and provides a proxy for spotted lanternfly adult and nymph density at our sites (sampled in summer 2019 – Table S1, Fig S2, Supporting Information 1, section 3). To test the effects of the explanatory variables on the number of eggs per egg mass, we fit a GLMM with a negative binomial error distribution (“glmer.nb” in `lme4`)with a nested random effect of tree ID within site.

## Results

### Survey Summaries

For the statewide survey, we examined egg mass presence/absence from 7,363 sites across the spotted lanternfly’s invaded range in Pennsylvania, 2,834 of which had egg masses present. For the core-to-edge survey, during egg collection year 2018, we surveyed 52 sites, 12 of which had ≥ 1 egg mass present at the site and tree-scale data (DBH, species) recorded. Across these 12 sites, we collected 183 egg masses from 40 trees (8 species, 6 genera) with an average of 27.52 eggs per egg mass (range 1-95 eggs). For egg collection year 2019, 38 sites were surveyed, 25 of which had ≥ 1 egg mass present and tree-scale data recorded. We collected egg masses at 19 of these 25 sites; we collected 280 egg masses from 81 trees (12 species, 8 genera) with an average of 32.84 eggs per egg mass (range: 2-102 eggs).

### Landscape scale

At the landscape scale, oviposition was higher at sites with older establishment age (*P* < 0.001) and higher at sites with low mean canopy cover surrounding the site (*P* < 0.05, Table 1). Across the variables tested, collection year had the strongest effect, reflecting an overall higher probability of oviposition at sites in the 2019 than 2018 season (*P* < 0.001). The effect of these explanatory variables was estimated after accounting for positive spatial autocorrelation among sites (*P* < 0.001).

**Table 1.** Results from each model of oviposition habitat use at different spatial scales

| Model Scale | Response Variable | R<sup>2</sup> | Term | Estimate | Std. Error | Statistic | P-value |
| --- | --- | --- | --- | --- | --- | --- | --- |
| Landscape | Presence/absence egg masses at sites | 0.200 | (Intercept) | -2.521 | 0.108 | -23.457 | <0.001 |
|  |  |  | Establishment Age | 0.229 | 0.035 | 6.465 | <0.001 |
|  |  |  | Canopy Cover | -0.142 | 0.059 | -2.412 | <0.05 |
|  |  |  | Collection Year (2019) | 2.049 | 0.061 | 33.439 | <0.001 |
|  |  |  | Spatial Autocovariate | 0.978 | 0.071 | 13.753 | <0.001 |
| Site | Presence/absence egg masses on trees | 0.489 | (Intercept) | 9.674 | 2.256 | 4.283 | <0.001 |
|  |  |  | Establishment Age | -1.510 | 0.545 | -2.770 | <0.01 |
|  |  |  | Canopy Cover | -6.864 | 1.974 | -3.477 | <0.001 |
|  |  |  | Tree DBH | -0.206 | 0.350 | -0.588 | 0.556 |
|  |  |  | *Acer* | 2.380 | 0.432 | -5.507 | <0.001 |
| Site | Abundance of egg masses on trees | 0.365 | (Intercept) | 6.379 | 0.985 | 6.478 | <0.001 |
|  |  |  | Establishment Age | -0.755 | 0.321 | -2.348 | <0.05 |
|  |  |  | Canopy Cover | -2.744 | 0.920 | -2.984 | <0.01 |
|  |  |  | Collection Year (2019) | -0.280 | 0.288 | -0.974 | 0.330 |
|  |  |  | Tree DBH | -0.074 | 0.419 | -0.177 | 0.859 |
|  |  |  | *Acer* | 1.482 | 0.286 | -5.189 | <0.001 |
| Tree | Presence/absence of egg masses in tree tops | 0.443 | (Intercept) | -4.234 | 0.859 | -4.932 | <0.001 |
|  |  |  | Establishment Age | 0.006 | 0.369 | 0.016 | 0.987 |
|  |  |  | Collection Year (2019) | 0.640 | 0.586 | 1.091 | 0.275 |
|  |  |  | Tree DBH | 5.062 | 1.092 | 4.637 | <0.001 |
| Egg Mass | Number of eggs per egg mass | 0.089 | (Intercept) | 3.383 | 0.136 | 24.950 | <0.001 |
|  |  |  | Establishment Age | -0.045 | 0.040 | -1.122 | 0.262 |
|  |  |  | Canopy Cover | 0.011 | 0.111 | 0.097 | 0.923 |
|  |  |  | Collection Year (2019) | 0.196 | 0.052 | 3.771 | <0.001 |
|  |  |  | *Acer* | 0.046 | 0.057 | -0.806 | 0.420 |
|  |  |  | Egg Mass Density | -0.035 | 0.036 | -0.986 | 0.324 |

### Site scale

At the site-scale, we found qualitatively similar patterns for the two models exploring the effects of the explanatory variables on egg mass presence and abundance, with stronger effects on egg mass presence. Site-level canopy cover had the strongest effect in the models, with higher egg mass presence (*P* < 0.001) and abundance (*P* < 0.01) at sites with low canopy cover (Table 1). The tree-level variable *Acer*/Not *Acer* was the next strongest variable with higher egg presence (*P* < 0.001) and abundance (*P* < 0.001) on trees from the *Acer* genus. Finally, site-level establishment age had the third-strongest effect such that egg mass presence (*P* < 0.01) and abundance (*P* < 0.05) on trees was higher at more recently invaded sites with lower establishment ages. This means that site-level establishment age has opposite effect on oviposition habitat use when considered at the site (negative effect size) and landscape scales (positive effect size).

### Tree scale

At the tree-scale, only tree DBH explained the likelihood to oviposit in the treetops versus in the entire tree, with trees with larger DBH being more likely to have eggs laid only in the treetop (*P* < 0.001, Table 1).

### Egg mass scale (fitness)

At the egg mass-scale, no site-level or tree-level variables explained variation in numbers of eggs per egg mass. The number of eggs per mass was only affected by collection year with significantly higher numbers of eggs per mass in collection year 2019 (*P* < 0.001, Table 1). There was also no evidence for density-dependent effects on fecundity.

## Discussion

Using a hierarchical, multiscale design, we identified several habitat associations for spotted lanternflies at the landscape, site, and tree scales which have important implications for spotted lanternfly management strategies. In addition, we found that the duration spotted lanternflies have been established at a site was also a predictor of habitat use, regardless of the habitat type. Remarkably, none of the environmental variables from the habitat associations nor the establishment age of sites explained variation in spotted lanternfly fecundity, indicating more work is needed to identify drivers of variation in spotted lanternfly fitness.

Across hierarchical scales of oviposition habitat use, the strongest explanatory power was at the broadest scales – landscape scale (oviposition at sites) and site-scale (oviposition on trees within sites). Oviposition at both scales was explained in part by the site-level explanatory variables, canopy cover and establishment age. Oviposition was associated with lower canopy cover in the 500m radius surrounding the site at both scales and is consistent with our hypothesis that spotted lanternfly oviposition is associated with human-dominated habitats. While the correlative nature of our study does not indicate a clear mechanism, the association with lower canopy cover habitats could be driven by selecting habitats that provide high fitness due to the presence of host plants and/or dispersal.

In the context of selecting habitats that provide high fitness, oviposition habitat selection in insects may be due to females choosing oviposition habitats that best guarantee offspring success, or may be a consequence of females choosing a habitat that is best for them at the time of oviposition (Mayhew, 1997). Although we cannot distinguish between these two oviposition habitat selection patterns for spotted lanternfly based on the influence of canopy cover alone, the higher oviposition on *Acer* genus trees we identified in the site-scale analyses may suggest oviposition habitat is based on adult preferences rather than selecting ideal host plants for offspring. *Acer* spp. phenology aligns better with the adult than nymph stage for feeding. Early instar nymphs preferentially feed on soft tissue such as herbaceous plants and fleshy parts of woody plants (Mason et al., 2020). While nymphs can feed on fleshy leaves of *Acer* species, the nymphs hatch in the spring often before leaf-out in most *Acer* species. Comparatively, adult lanternflies frequently congregate on *Acer* spp. in autumn (Mason et al., 2020), likely as a consequence of the delayed autumnal senescence and prolonged photosynthetic activity that characterizes members of the *Acer* genus relative to other regional tree species. Alternatively, oviposition on *Acer* spp. may reflect its utility as an oviposition substrate rather than a trophic resource due to its relatively smooth bark which may be a preferred oviposition substrate (Urban, 2020). Like *Acer* spp., *Ailanthus altissima* also has smooth bark and delayed phenology, was second to *Acer* in our study as an oviposition substrate and has been used as an oviposition substrate in other locations within the invaded range (Liu, 2019). Future experimental work should aim to disentangle the effects of phenology from substrate properties on oviposition substrate selection mechanisms.

In addition to spotted lanternfly possibly selecting low canopy cover habitats due to presence of host plants, the association between higher oviposition and lower surrounding canopy cover may also reflect spotted lanternfly’s natural and/or human-assisted dispersal. Prior to oviposition, lanternfly adults disperse by flying. As their dispersal capabilities are limited in flight length and distance (Wolfin et al., 2019), it may be easier for them to disperse in areas with less dense canopy cover. This would result in a positive association between oviposition and lower canopy cover like we observed in our data. Additionally, all life stages of spotted lanternfly are transported short distances within the invaded range by humans (Urban et al., 2021). Human-assisted dispersal would likely result in them being more tightly associated with human-impacted (i.e., low canopy cover) areas.

Compared to the consistent effect of canopy cover on oviposition at the landscape and site scales, site-level establishment age had opposing effects at these two scales. These opposing effects indicate that sites that have been invaded longer are more likely to have at least some egg masses present, yet individual trees at sites invaded more recently are more likely to have egg masses present and a higher abundance of egg masses. Although studies of spotted lanternfly population dynamics are ongoing, they appear to match classic invasion dynamics where newly colonized sites experience a rapid increase in population density and then decline to low densities without ever reaching a point of extinction at the site (Burton et al., 2010). This boom-and-bust population dynamics pattern supports our finding that older sites are more likely to have eggs present because extinction is rare at old sites, and colonization of sites at the invasion edge is patchy. In comparison, our site-scale results suggest that at sites with younger establishment age, spotted lanternflies may have a higher likelihood of ovipositing on a range of trees at a site rather than concentrating their eggs on a few trees, regardless of the tree species. Whether ovipositing on a range of trees occurs due to spillover after the prime sites on preferred trees are occupied when population densities are high or is a pattern that happens independent of population densities is currently unclear.

Collection year also explained significant variation in oviposition with a higher likelihood of oviposition at sites in 2019 (landscape scale) and more eggs per egg mass in 2019 (egg mass scale). While the exact mechanism behind annual variation in reproductive output in spotted lanternflies is not known, annual temperature fluctuations may be responsible for the patterns seen. For example, slight elevations in warming (i.e, 2°C) can increase the number of eggs laid per female in other planthoppers (Manikandan et al., 2015). The yearly average temperature was warmer and precipitation was lower in 2019 compared to 2018 in southeastern Pennsylvania (NCEI, 2019, 2020), so this may have contributed to the seasonal fluctuations we documented.

Aside from collection year, no other variables we tested explained significant variation in spotted lanternfly fecundity despite there being a substantial range in the number of eggs per egg mass in our data (1-102 eggs). Lanternfly reproductive biology is currently under investigation including determining the number of egg masses each female lays and the allocation of resources to egg masses. In insects, habitat quality can affect reproductive output beyond the number of eggs per egg mass, including the size of eggs, occurrence of offspring deformities, and offspring survival (Awmack & Leather, 2002). In addition, insects can adjust egg quality and nutrient allocation to eggs to match environmental conditions and lay higher quality eggs in better habitats (Awmack & Leather, 2002). There was no influence of intraspecific competition (lanternfly density), human-dominated habitats (represented by canopy cover), or establishment age on the number of eggs per mass in our study. Clearly more research is needed to understand variation in lanternfly fecundity and to determine whether other aspects of reproductive output are useful metrics for fitness.

The multiscale nature of our results suggest explicit management strategies are needed at both the site and tree scales to locate egg masses, in addition to differing strategies at recently established sites at the invasion edge versus long-established sites within the invasion core (Fig. 3A). For sites, our data suggest higher variation in the presence of egg masses at younger sites at the invasion edge relative to the core (Fig. 3B). At the tree scale, if eggs are present at a site, trees at younger sites are more likely to have egg masses than older sites (Fig 3B). In practice, this means older sites are more likely to have egg masses present, but they may be at lower density and possibly harder to find. Comparatively, egg mass densities are likely to be larger and distributed across more trees at younger sites. Across the invasion gradient, we emphasize the need to treat possibly overlooked *Acer* spp., especially in greenspaces and residential areas where *Acer rubrum* is frequently used as an ornamental tree (Mason et al., 2020). We caution that egg masses in larger trees may be inaccessible for mechanical removal because they may be in the tree canopy and other methods may be necessary (Leach et al., 2019). Finally, our results have important implications for optimal control strategies for invasive pests in general. Recent theoretical work indicates that although controlling the populations that are the source of propagules is optimal, whether those source populations are more at the edge or the core of the invasion may be species-specific (Baker, 2017). Our results suggest different search methods may be needed depending on whether core or edge populations are targeted for control.

**Figure 3:**
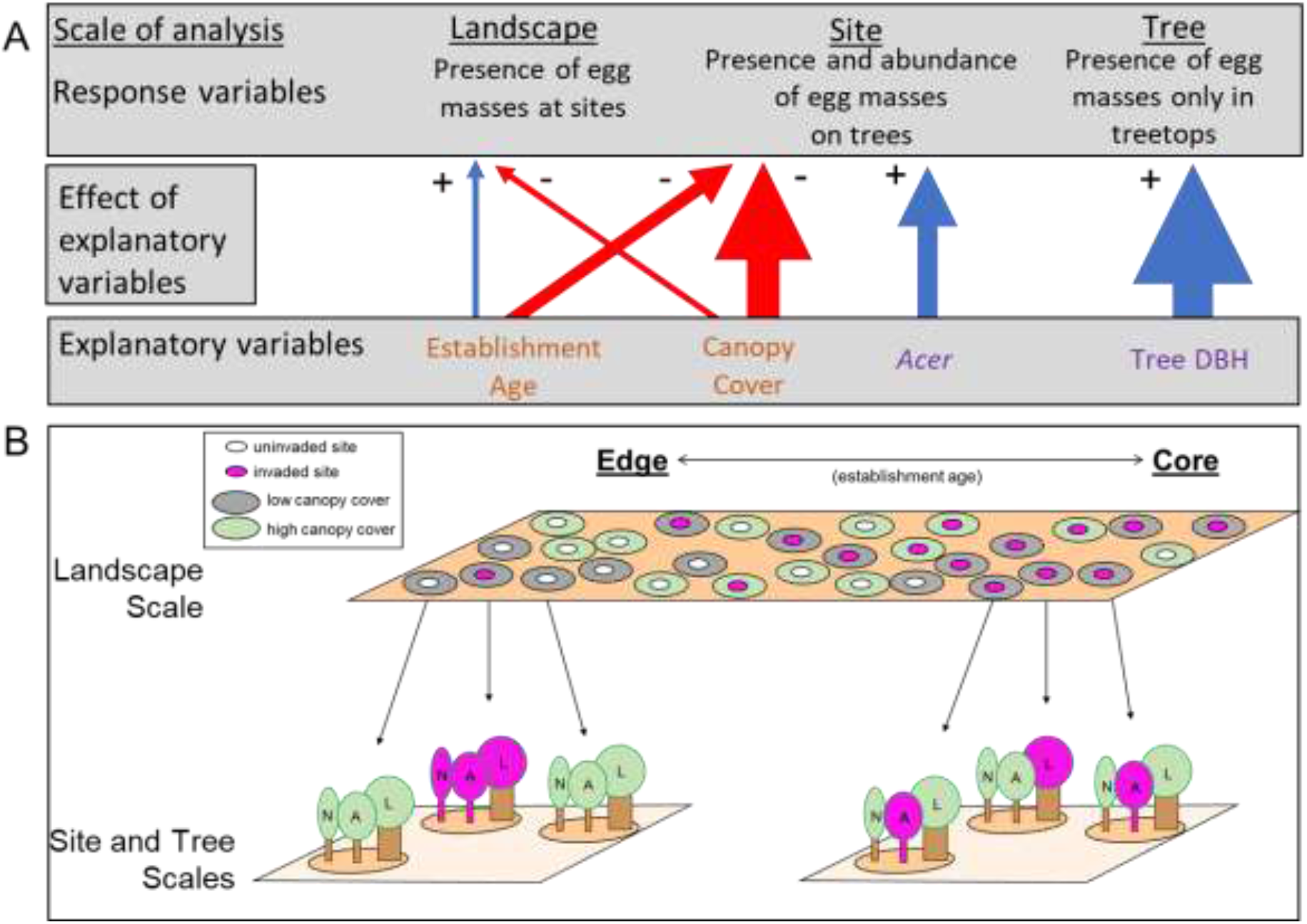
A) Main effects from multiscale models that inform management strategies. Width of arrow is proportional to effect size; blue arrows indicate positive effect, red arrows are negative effect of site-level (orange) or tree-level (purple) explanatory variables on multiscale response variables. B) Pattern of spotted lanternfly oviposition habitat associations (pink shading signifies lanternfly oviposition habitat use) at the landscape and site/tree scales that indicates different search strategies are needed to control spotted lanternfly along the gradient of establishment ages for sites. At the landscape scale, more sites at the core will have eggs present. When eggs are present at a site, more trees at sites at the edge will have egg masses than at the core. Across the gradient, sites with low canopy cover in the surrounding landscape (gray circles) and trees from the *Acer* genus (A) versus other genera (N) will have a higher likelihood of having spotted lanternfly egg masses. In addition, large trees (L) will have a higher likelihood of having egg masses at the top of the trees that are out of reach.

## Author Contributions

VAR, MRH and JEB conceived the ideas and designed the methodology; VAR collected the data; VAR and SDB led the analyses with input from MRH and JEB; VAR and JEB led the writing of the manuscript. All authors contributed critically to the drafts and gave final approval for publication.

## Acknowledgements

We thank Dawn Becker, Adam Gasiewski, Ben Gress, Renee Johnson, and Jeffery Stuart who helped with data collection; Nadège Bélouard, Stephanie Lewkiewicz, and Benni Seibold for feedback on the manuscript; and the iEcoLab at Temple University for helpful discussions. This work was funded by the USDA Animal and Plant Health Inspection Service Plant Protection and Quarantine (AP19PPQS&T00C251, AP20PPQS&T00C136); USDA SCRI (2019-51181-30014); the PA Department of Agriculture (44176768, 44187342, C9400000036, C94000833); and a Francis Velay fellowship awarded to VAR. This research includes calculations carried out on HPC resources supported in part by the National Science Foundation through major research instrumentation grant number 1625061 and by the US Army Research Laboratory under contract number W911NF-16-2-0189.

## Data Availability Statement

Data DOI will be made available upon acceptance.

## Conflict of Interest

The authors declare no conflict of interest.

## Notes

### Competing Interest Statement

The authors have declared no competing interest.

