## Supplement for "Multiscale assessment of oviposition habitat associations and implications for management in the spotted lanternfly (*Lycorma delicatula*), an emerging invasive pest"

**Supporting Information 1**

*Section 1. Calculating establishment age through interpolation*

Our goal was to estimate the duration each site in our dataset had an established population of spotted lanternfly. We spatially interpolated the presence/absence of spotted lanternfly at any life stage from data collected by PDA and USDA across PA for each year between 2015 and 2019. To do the interpolation, we aggregated data at a 1 km grid scale, considering each grid as positive for spotted lanternfly if at least one point within the cell was recorded as having an established population. If a cell was scored as positive for spotted lanternfly in a given year, it was considered positive thereafter. We then used inverse distance weighing (with a weighing factor of 4) to estimate the probability of spotted lanternfly being established in a site in a given year as a value between 0 (certainly absent) and 1 (certainly present). Finally, we calculated the establishment age of a site as a continuous variable by combining the probabilities of spotted lanternfly being established across the years (2015 to 2019). The maximum value for any site was 5 years indicating certainty of spotted lanternfly being established throughout the 5 years considered, between 2015-2019 (Fig. 2). Given that the spatial coverage of the statewide survey encompassed our core-to-edge survey sites, this interpolation method allowed us to calculate establishment ages for all study sites from both survey data sets that we could then use in analyses at all scales.

*Section 2. Tree Genus Oviposition Preference*

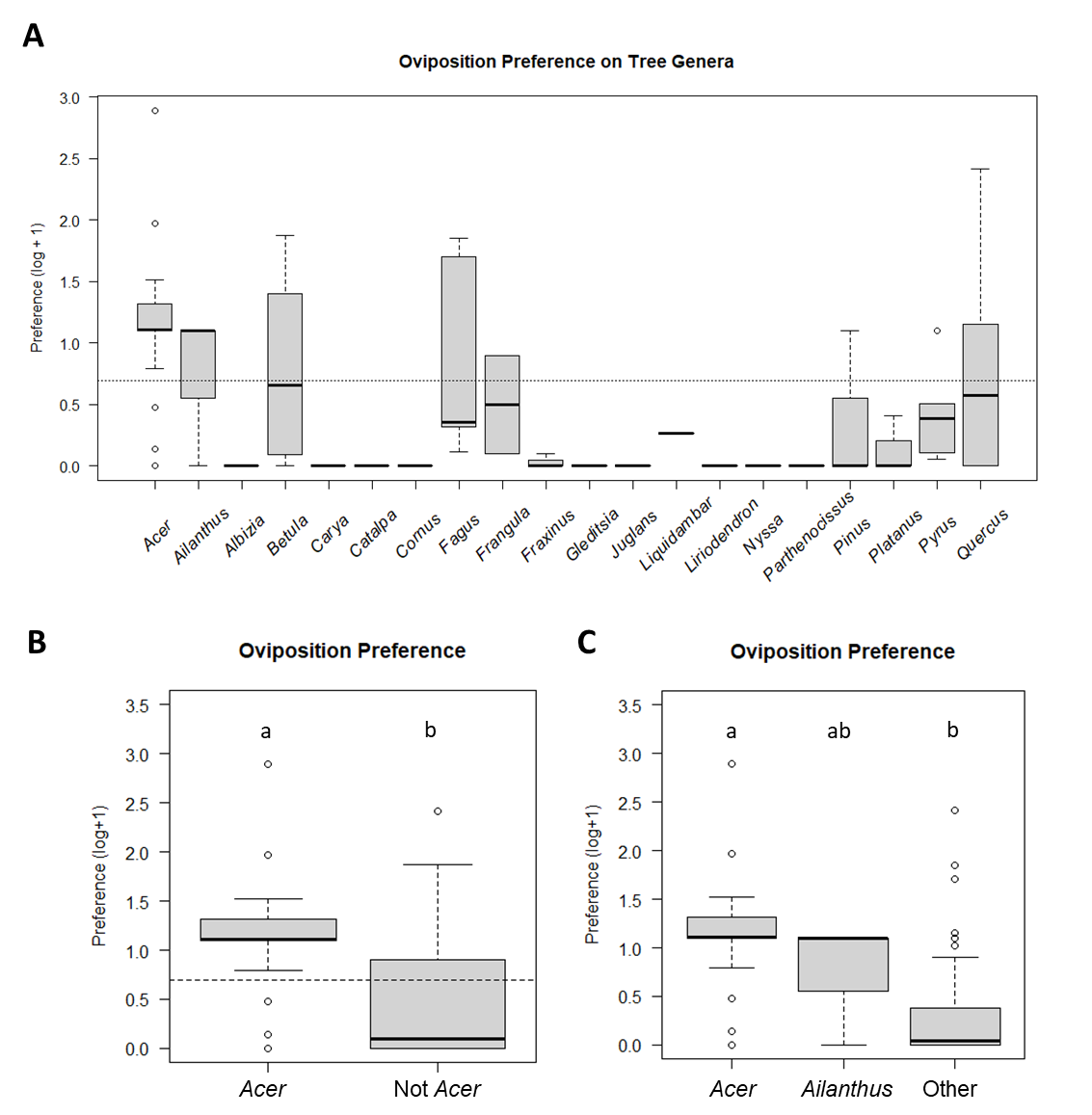

**Figure S1:** Evidence for oviposition preference for tree genera. Preference was measured as the proportion of egg masses on a genus at a site relative to the proportion of total available substrate for a genus (tree DBH) at a site. Horizontal dotted lines indicate oviposition selection on tree genera is exactly proportional to the abundance of that genera at sites (i.e. log[(preference=1)+1]). **A.** There was a significant effect of tree genus on oviposition preference (*F*_18,64_=2.65, *P*<0.01). **B.** When we explored this pattern, we found a significantly higher preference for genus *Acer* relative to all other genera combined (*F*_1,83_=29.89, *P*<0.001). **C.** After *Acer*, spotted lanternflies exhibited the next strongest preference for *Ailanthus.* However, the preference for *Ailanthus* did not differ significantly from *Acer* (*P* = 0.44) or the rest of the genera (*P* = 0.47*)*, therefore we used the binary *Acer*/not *Acer* variable in our analyses going forward. In panels B and C, lower means with the same lower-case letter are not statistically different at *P*<0.05.

*Section 3. Link between egg mass density and SLF density*

Beginning at the start of hatch season on May 20, 2019, we returned to the same sites that were surveyed in collection year 2018 to sample spotted lanternfly densities on trees using sticky band traps. We banded trees where eggs were collected in collection year 2018 as well as additional trees within a 10m radius of the focal trees. Bands were left on trees for 24h, after which we returned to sites remove bands and count the number of lanternflies per instar on each band. Each site was sampled 3-4 times during the season to try and sample each instar, with the final sampling occurring on September 11, 2019. During the 2019 collection year, we again returned to these sites to survey for and collect egg masses.

We regressed the number of egg masses recorded on trees at each site during the 2019-20 collection year on the number of lanternflies collected on sticky bands to determine if egg mass density was a reasonable reflection of live spotted lanternfly density. We found a positive relationship between the density of spotted lanternfly on sticky bands and egg masses recorded on trees (R^2^=0.99, *P* < 0.001; Table S1; Fig. S2).

**Figure S2**: The number of egg masses observed on trees is representative of adult and nymph lanternfly densities in the previous summer recorded on sticky band traps at 23 sites across southeastern Pennsylvania.

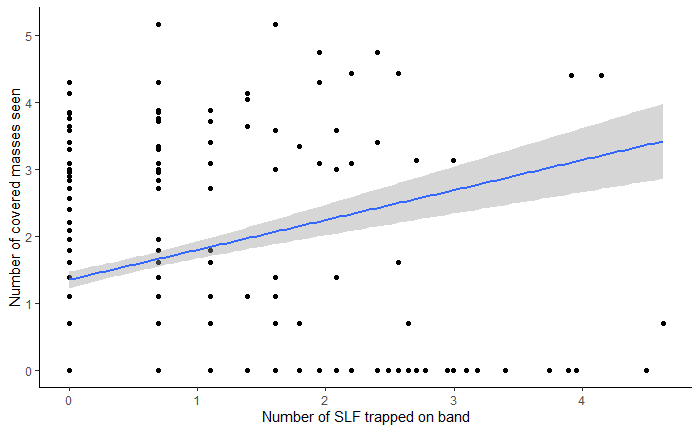

**Table S1:** Results from the model fitting the number of egg masses to the number of SLF trapped on a sticky band at each tree

| **Term** | **Estimate** | **Std. Error** | **Statistic** | **P-.value** |
| --- | --- | --- | --- | --- |
| Intercept | 0.049 | 0.650 | 0.075 | 0.940 |
| Number of SLF trapped on band | 0.241 | 0.009 | 26.343 | **<0.001** |
